# The Influence of Variable-Heavy (V_H_) Chain Families on IgG_2_, _3_, _4_ on FcγRs and Antibody Superantigens Protein G and L Binding using Biolayer Interferometry

**DOI:** 10.1101/2023.03.26.534243

**Authors:** Anthony M. Deacy, Samuel Ken-En Gan

## Abstract

**Background:** As the most abundant immunoglobulin in blood and the most common human isotype used for therapeutic monoclonal antibodies, the engagement and subsequent activation of its Fc receptors by IgGs are crucial for antibody function. While generally assumed to be relatively constant within subtypes, recent studies have shown the antibody variable regions to exert distal effects of modulating antibody–receptor interactions on many antibody isotypes. Such effects are also expected for IgG and its subtypes with the in-depth understanding of these V-region effects highly relevant for engineering antibodies, antibody purifications, and understanding to how robust the microbial immune evasion proteins are.

**Methods:** In this study, we created a panel of IgG_2_/IgG_3_/IgG_4_ antibodies by changing the V_H_ family (V_H_1-7) frameworks while retaining the complementarity determining regions of Pertuzumab and measured the interaction of the IgGs with FcγRIa, FcγRIIa_H167_, FcγRIIa_R167_, FcγRIIb/c, FcγRIIIa_F176_, FcγRIIIa_V176_, FcγRIIIb_NA1_, and FcγRIIIb_NA2_ receptors alongside antibody superantigens proteins L and G using biolayer interferometry.

**Results:** The library of 21 IgGs demonstrated that the V_H_ frameworks influenced receptor binding sites on the constant region of the subtypes significantly, providing non-canonical interactions and non-interactions. However, there was minimal influence on the binding of bacterial B-cell superantigens Proteins L and G on the IgGs, showing their robustness against V-region effects.

**Conclusions:** These results demonstrate the importance of the V-regions during humanization of therapeutic antibodies that can confer or diminish FcR-dependent immune responses, while remaining both suitable and susceptible to the binding by bacterial antibody superantigens in antibody purification and be present with normal flora.

**STATEMENT OF SIGNIFICANCE:** IgGs are the predominant isotype for clinical and research applications. Despite the vast amount of research to study it, particularly on IgG1, there remains a gap in understanding how the variable regions and the receptor binding sites can influence one another in the other IgG subtypes, across the IgG subtypes with different hinges and makeup. This study investigates the effect of these variable regions on the engagement of receptors and also how bacterial antibody superantigens present in microflora and used in antibody purification can exert distal effects.

## INTRODUCTION

IgG is the most common immunoglobulin found in human blood (10 – 20 %) [1] and is the predominant antibody isotype used in research and clinically. Numerous studies on the latter clinical applications and characterization focus on its safety, selectivity, diversity, solubility, tolerability, stability, and half-life [2]. Yet, leveraging on antibody constant regions to confer localization [2, 3], reducing systemic circulation to mitigate side effects, lower dosages [4] and other functions often remain neglected in characterization. A possible reason for the application gap is the concern about unexpected effects from other antibody regions. Considering previous findings on IgG_1_ where the areas of V_H_-V_L_ affected FcγRIIa binding at the IgG_1_ heavy chain constant (C_H_), that was followed by similar findings on other isotypes: IgE [5], IgA_1_ and _2_ [6, 7] and on IgM [8], only secretory IgD and the rest of the IgG subtypes remain to be characterized.

Antigen binding by IgG_1_ and IgG_2_ can increase the binding affinity between the antibody and its FcγR [9] from effects originating from the V-regions. Such allosteric effects were contributed by both the complementarity-determining-regions (CDRs) and frameworks (FWRs) of the heavy chain, as well as the variable light chain (V_L_) FWR [10]. This effects were also observed in the antibody–antigen interaction for IgG_4_ when introducing mutations several nanometres away [11], indicating the need to elucidate how mutations outside the CDRs in the V-FWRs can influence the FcγR binding sites.

The human IgG is further categorized into four subclasses or subtypes: IgG_1_ (60 – 70 % total IgG), IgG_2_ (20 – 30 % total IgG), IgG_3_ (5 – 8 % total IgG) and IgG_4_ (∼5 % total IgG) [12]. This classification is based on the C_H_ chain, which shares ∼90% homology among the subtypes. Yet, the differences can result in significant variation at the antibody hinge regions and the engagement to immune complement proteins and FcγRs [1, 13]. For human IgGs, there are nine common Fragment crystallizable γ receptors (FcγRs) (FcγRIa, FcγRIIa_H167_, FcγRIIa_R167_, FcγRIIb/c, FcγRIIIa_F176_, FcγRIIIa_V176_, FcγRIIIb_NA1_, FcγRIIIb_NA2_ and FcRn), each showing different interactions with the various IgG subtypes [1, 14]. IgG_3_ was previously reported to bind more strongly to FcγRIIa, FcγRIIIa and FcγRIIIb than IgG_1_, and that IgG_2_ and IgG_4_ bound weakly to FcγRIIa, FcγRIIIa and FcγRIIIb [15].

FcγRIa is the only high-affinity FcγR expressed on the surface of dendritic cells, neutrophils, monocytes, macrophages, microglia, eosinophils, basophils mast cells and platelets [16]. Unlike other FcγRs, FcγRIa can be stimulated as a monomer. Activating the immunoreceptor tyrosine-based activation motif (ITAM) to result in antibody-dependent cellular phagocytosis (ADCP) and cytokine release [17]. FcγRIIa is sub-classified into FcγRIIa_H167_ and FcγRIIa_R167_. Both are low-affinity receptors that require multimerization for activation and are expressed in neutrophils, eosinophils, monocytes, macrophages, dendritic cells, platelets, basophils, microglia, mast cells and CD4^+^ T-cells [16]. Stimulation of FcγRIIa utilizes the ITAM pathway to induce antibody-dependent cellular cytotoxicity (ADCC) and ADCP [17].

FcγRIIb/c are low-affinity receptors expressed in neutrophils, eosinophils, monocytes, macrophages, dendritic cells, B cells, basophils, natural killer cells, microglia and mast cells [16] that multimerize during activation. FcγRIIb is the only direct inhibitory FcγR and, unlike other FcγR, activates the immunoreceptor tyrosine-based inhibition motif (ITIM) to inhibit ADCC, ADCP and B-cell activation [17]. Activation of FcγRIIc, on the contrary, results in the activation of ADCC, ADCP and B-cell activation via the ITAM signalling pathway [17].

FcγRIIIa is another low-affinity receptor, sub-classified into FcγRIIIa_F176_ and FcγRIIIa_V176_. It is expressed in monocytes, dendritic cells, macrophages, neutrophils, natural killer cells, monocytes, microglia, CD4^+^ T-cells and CD8^+^ T-cells [16], also using the ITAM signally pathway to activate ADCC and ADCP [17].

FcγRIIIb is a low-affinity inhibitory receptor sub-classified into FcγRIIIb_NA1_ and FcγRIIIb_NA2_ found in neutrophils, dendritic cells, neutrophils, monocytes, macrophages, eosinophils and basophils [16]. The activation of FcγRIIIb results in the inhibition of ADCP through its function as a decoy receptor [17].

Apart from binding FcR and the antigen, many Igs also interact specifically with many bacterial B-cell superantigens that bind at the V-region FWRs [18]. Bacterial B-cell superantigens are multidomain proteins that can activate the B-cell receptor (BCR) on the surface of B cells, leading to B-cell apoptosis [19]. Previous studies have shown bacterial B-cell superantigen (Protein A and Protein L) binding to remain relatively stable when changing the combination of V_H_ – V_L_ chains [20].

Since the V-regions can determine superantigen recognition and also FcR binding [20], having a combined investigation of these interactions is essential, especially given the recent interest in Fc engineering to modify antibody functions [21, 22]. This investigation thus aims to build on previous work on IgG_1_ to fill the gap by focusing on IgG_2_, IgG_3_ and IgG_4_ that were grafted with the CDRs of Pertuzumab. By varying the 7 V_H_ families, 21 antibodies were investigated with the FcγRs using biolayer interferometry to provide a foundational understanding of the superantigen-antibody-FcR interaction.

## MATERIALS AND METHODS

### Antibody Production

All Pertuzumab V_H_ sequences IgG_2_,_3_ and_4_ were recombinantly made as described previously [3] and joined with our previously Pertuzumab CDR grafted V_H_1-7 [23] to make the respective recombinant V_H_1-7 IgG_2-4_. Briefly, the inserts were PCR-amplified out with the adaptors and ligated, followed by transformation into competent *E. coli* (DH5α) strains [24, 25] and plasmid extraction [26] and transfected into HEK293 cells [3]. Transfection, production, and purification were performed previously [5, 27] using HEK293 cell lines transient transfection.

### Binding Affinity Quantification

Commercial Avi-tagged receptors: FcγRIa (Sino Biological Inc. Cat: 10256-H27H-B), FcγRIIa_H167_ (Acro Biosystems Cat: CDA-H82E6), FcγRIIa_R167_ (Acro Biosystems Cat: CDA-H82E7), FcγRIIb/c (Acro Biosystems Cat: CDB-H82E0), FcγRIIIa_F176_ (Acro Biosystems Cat: CDA-H82E8), FcγRIIIa_V176_ (Acro Biosystems Cat: CDA-H82E9), FcγRIIIb_NA1_ (Acro Biosystems Cat: CDB-H82E4) and FcγRIIIb_NA2_ (Sino Biological Inc. Cat: 11046-H27H-B) were immobilized onto a streptavidin biosensor (Sartorius Cat: 18-5020) and subjected to free-floating IgG variants to obtain the rate of association (K_a_), dissociation (K_d_) and equilibrium dissociation constant (K_D_). The program and steps used in the Octet Red ® system were described previously, along with the software calculation of the K_D_ [5-8, 23].

### BLI measurements of immobilized IgG receptors to Pertuzumab V_H_1-V_H_7 IgG_2, 3_ and _4_ variants

To measure the binding affinities of the different V_H_ -grafted Pertuzumab, the FcγRs were first immobilized to BLI streptavidin (SA) sensors via their Avi-tag with a minimum loading threshold target of 1 nM. The binding affinities of IgG_2_, IgG_3_ and IgG_4_ were measured using the Octet® Red 96 Biolayer Interferometry (BLI). The IgG subtypes _2_,_3,_ and _4_ were analyzed by representatives of the seven V_H_-FWR IgG_2,3,4_ with the Pertuzumab CDRs. The Pertuzumab binding affinity screen displayed a range of dissociation equilibrium constants (K_D_) from ∼0.06 nM to 0.5 μM (Tables 1, 2 and 3). All measurements were made in triplicates unless otherwise stated.

**Table 1:**
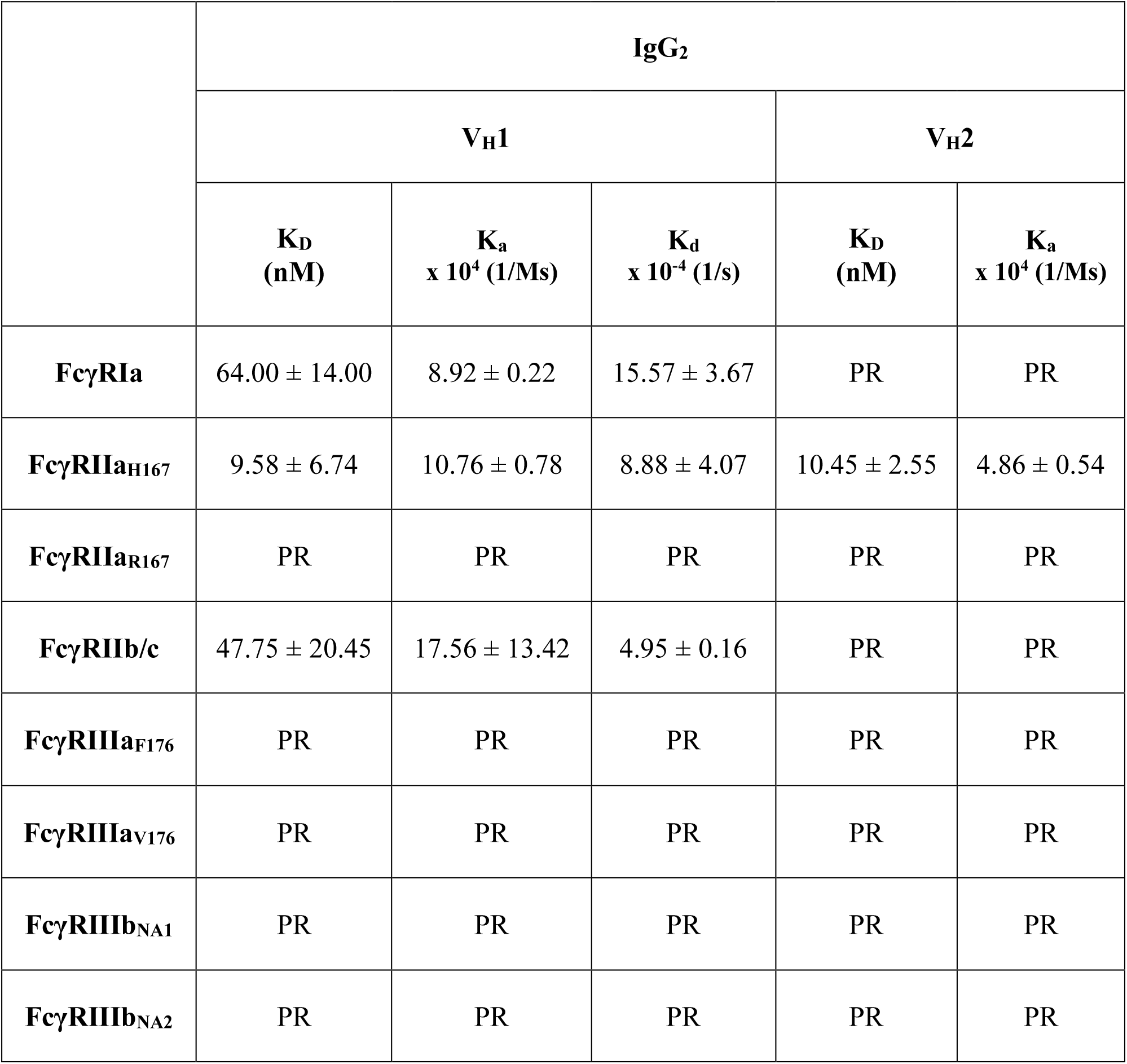
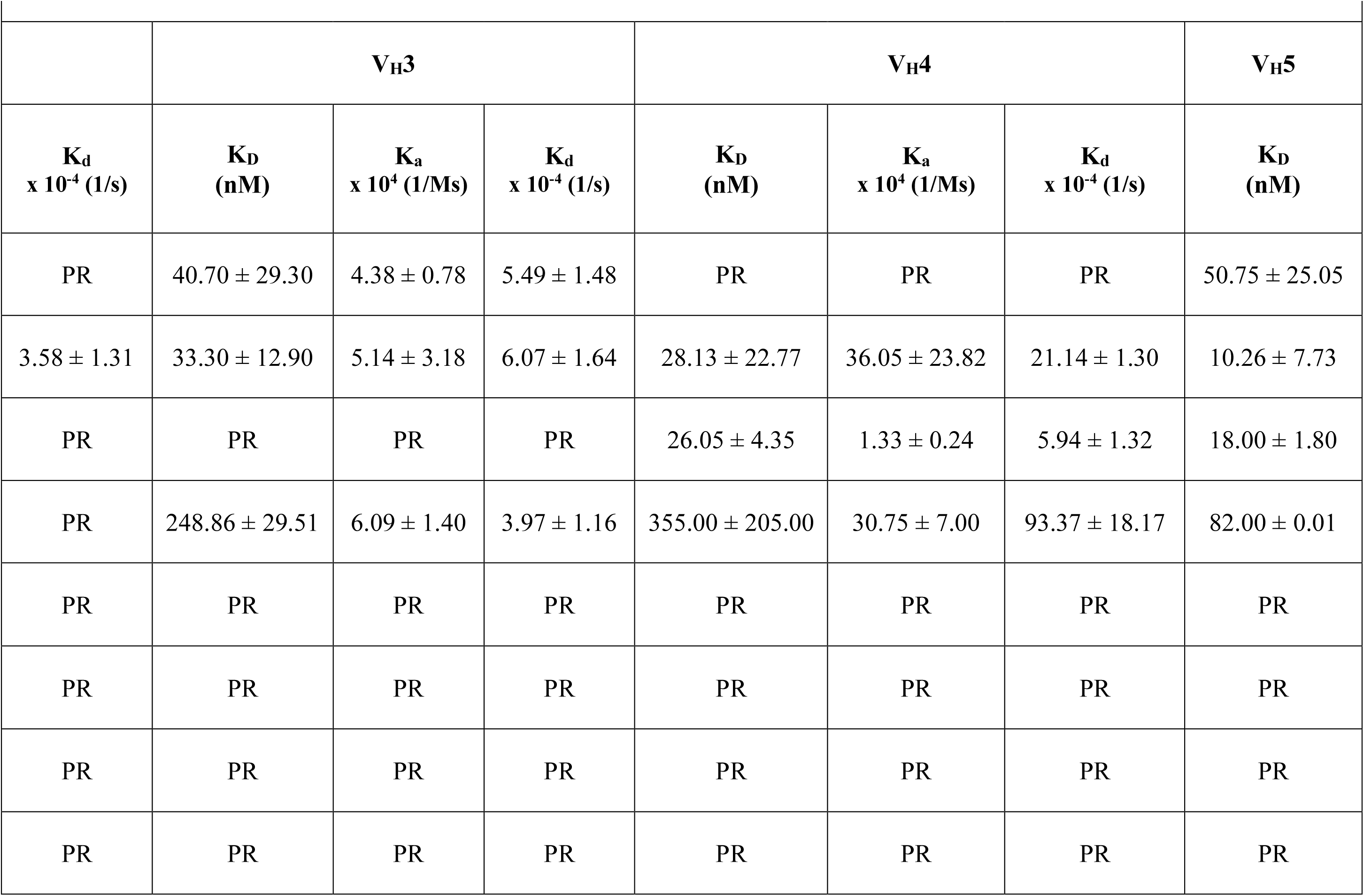
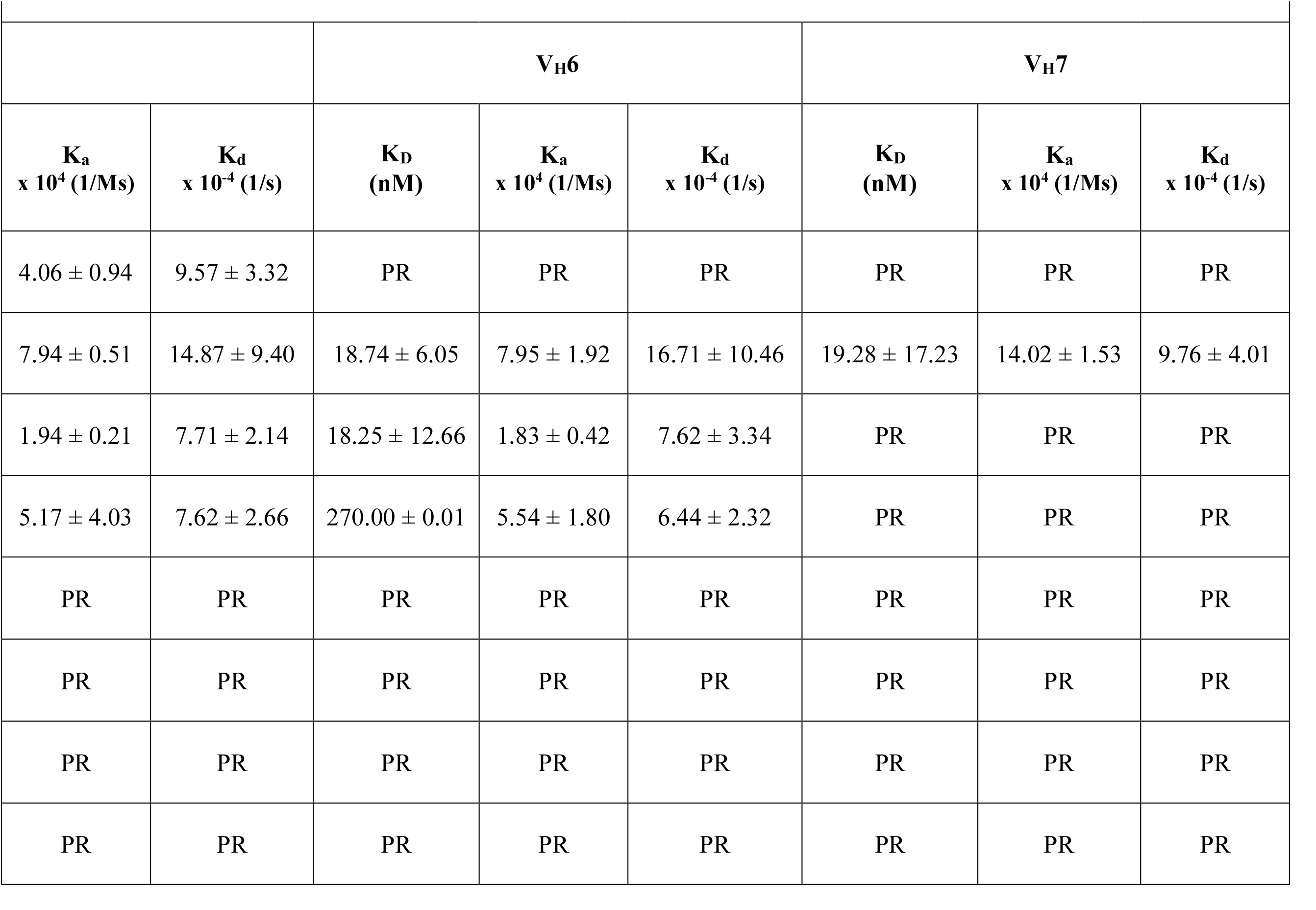
BLI measurement of Pertuzumab IgG_2_ V_H_1 to V_H_7 variants against immobilized FcγRs (FcγRIa, FcγRIIa_H167_, FcγRIIa_R167_, FcγRIIb/c, FcγRIIIa_F176_, FcγRIIIa_V176_, FcγRIIIb_NA1_ and FcγRIIIb_NA2_). K_D_ (nM), K_a_ (x 10^4^ (1/Ms)) and K_d_ (x 10^-4^ (1/s)) are all presented. PR indicates a poor response (RU below 0.3). All experiments were performed in triplicates with the standard errors shown.

**Table 2:**
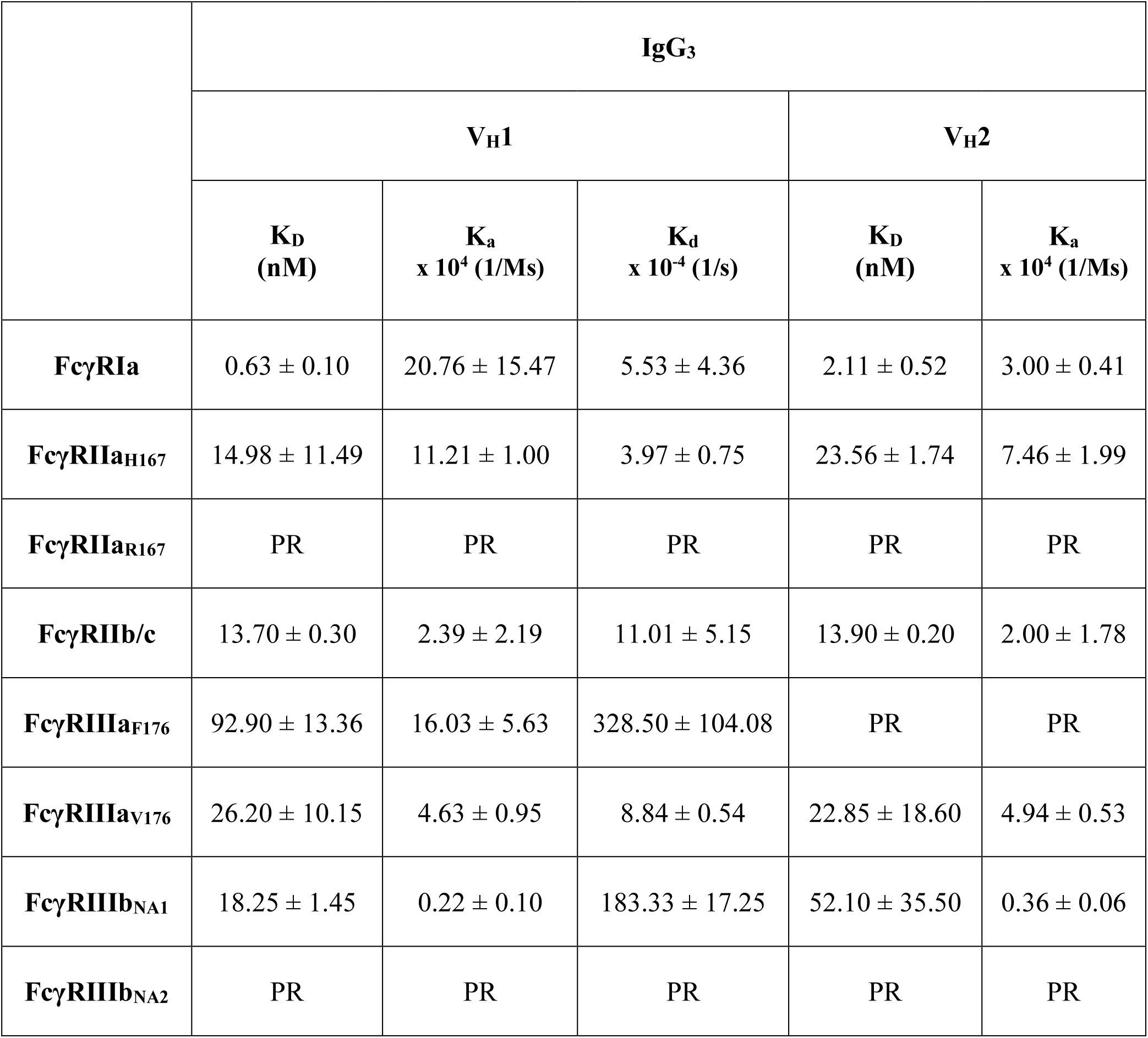
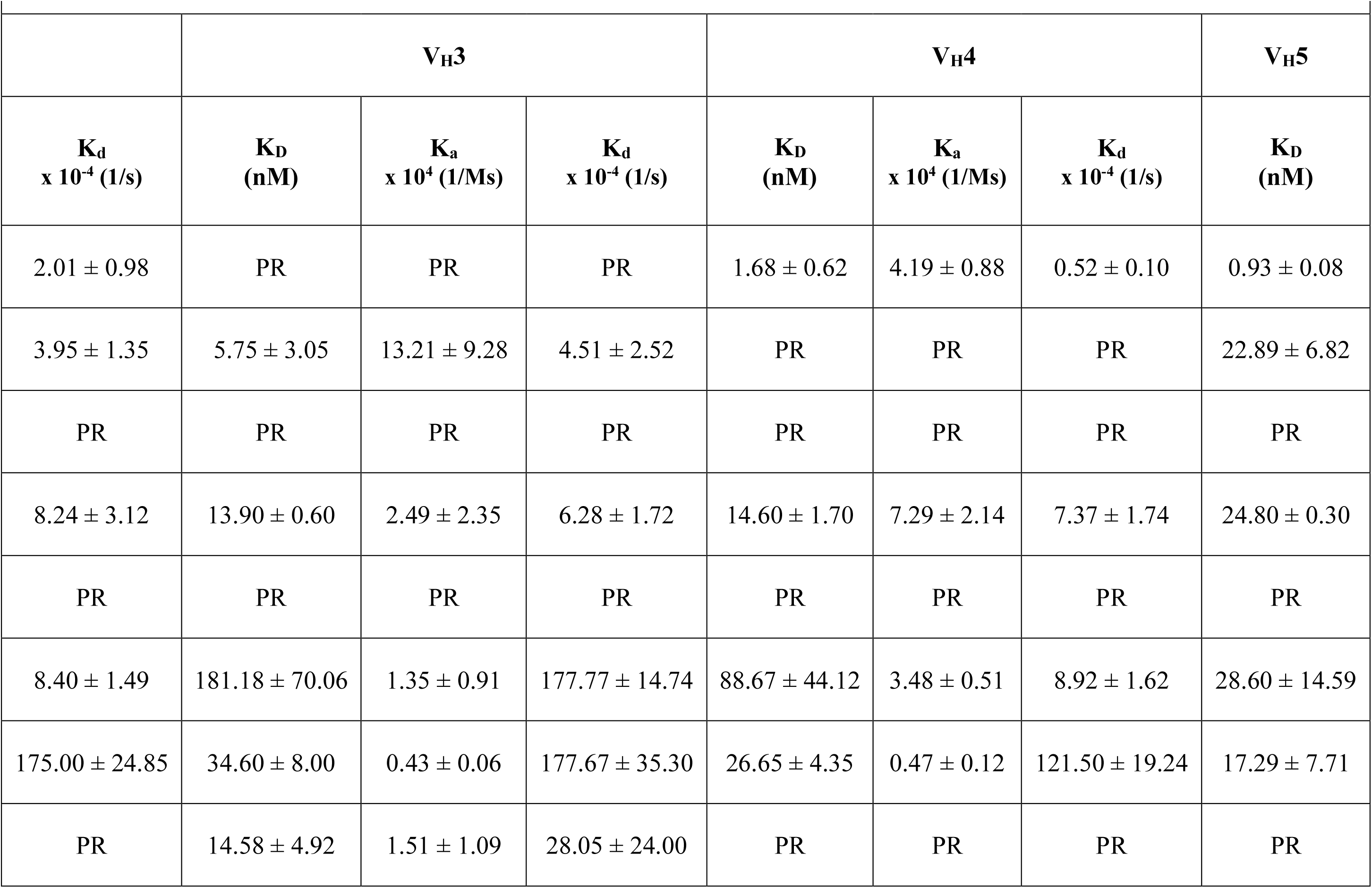
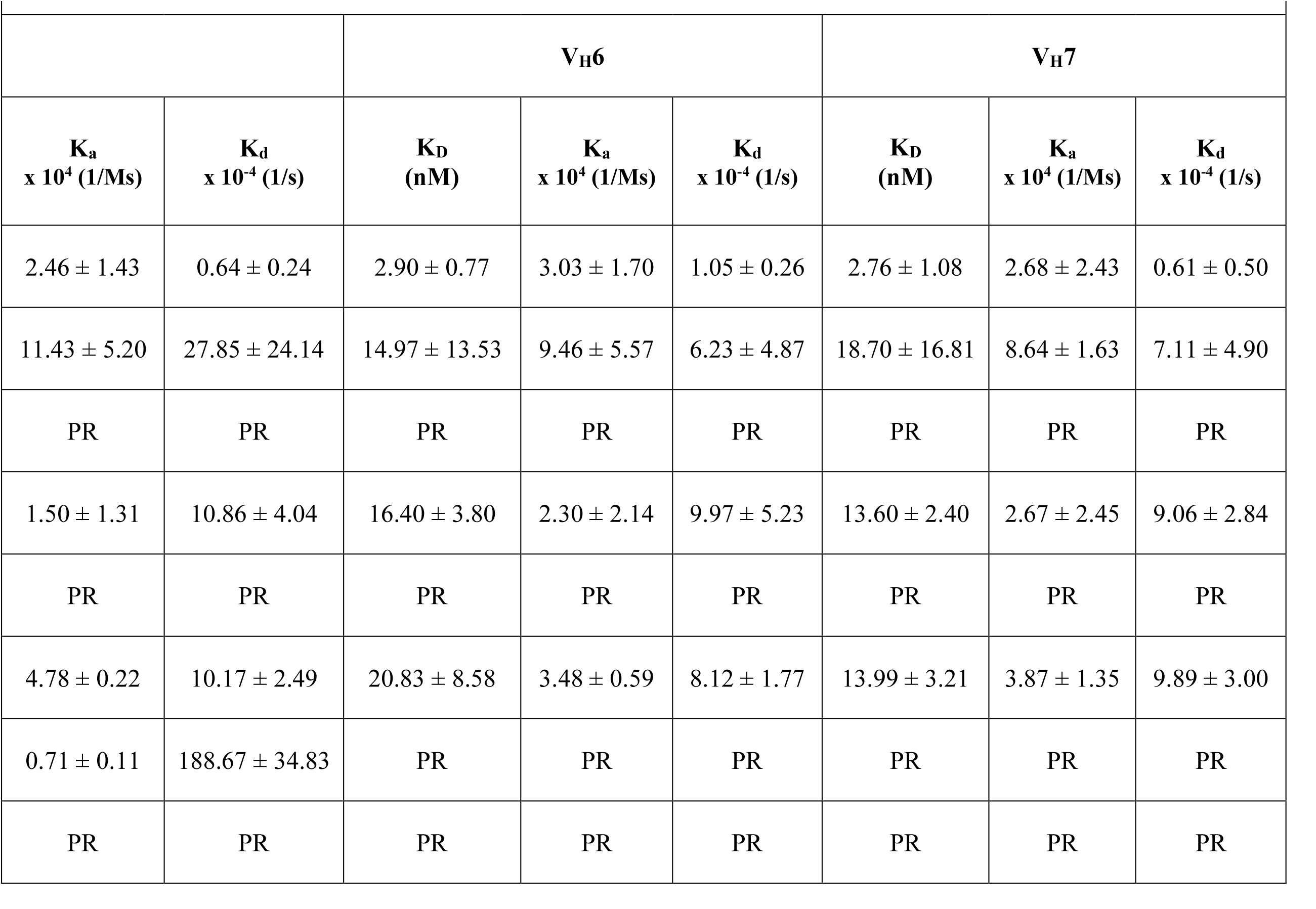
BLI measurement of Pertuzumab IgG_3_ V_H_1 to V_H_7 variants against immobilized FcγRs (FcγRIa, FcγRIIa_H167_, FcγRIIa_R167_, FcγRIIb/c, FcγRIIIa_F176_, FcγRIIIa_V176_, FcγRIIIb_NA1_ and FcγRIIIb_NA2_). K_D_ (nM), K_a_ (x 10^4^ (1/Ms)) and K_d_ (x 10^-4^ (1/s)) are all presented. PR indicates a poor response (RU below 0.3). All experiments were performed in triplicates, with the standard errors shown.

**Table 3:**
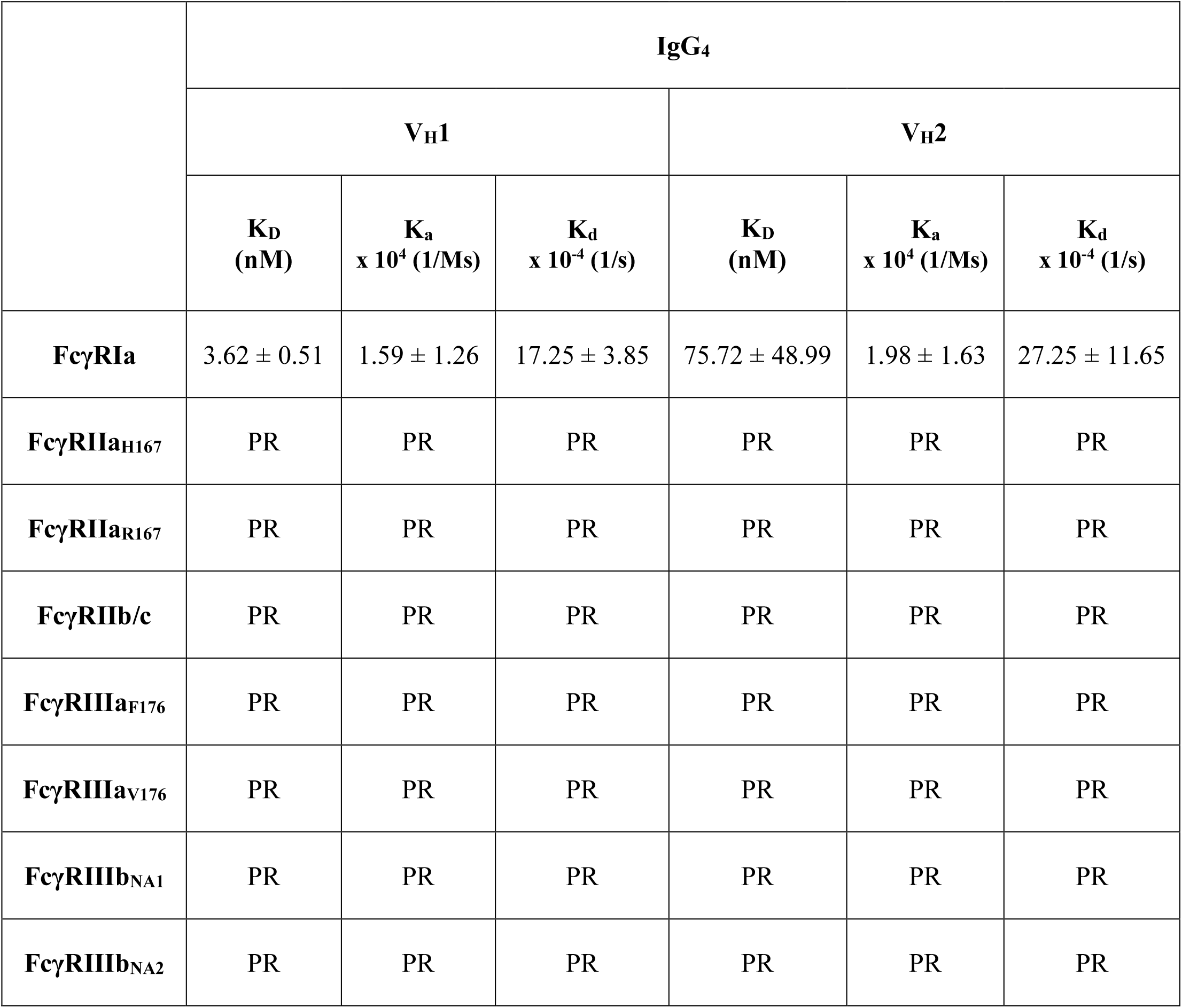
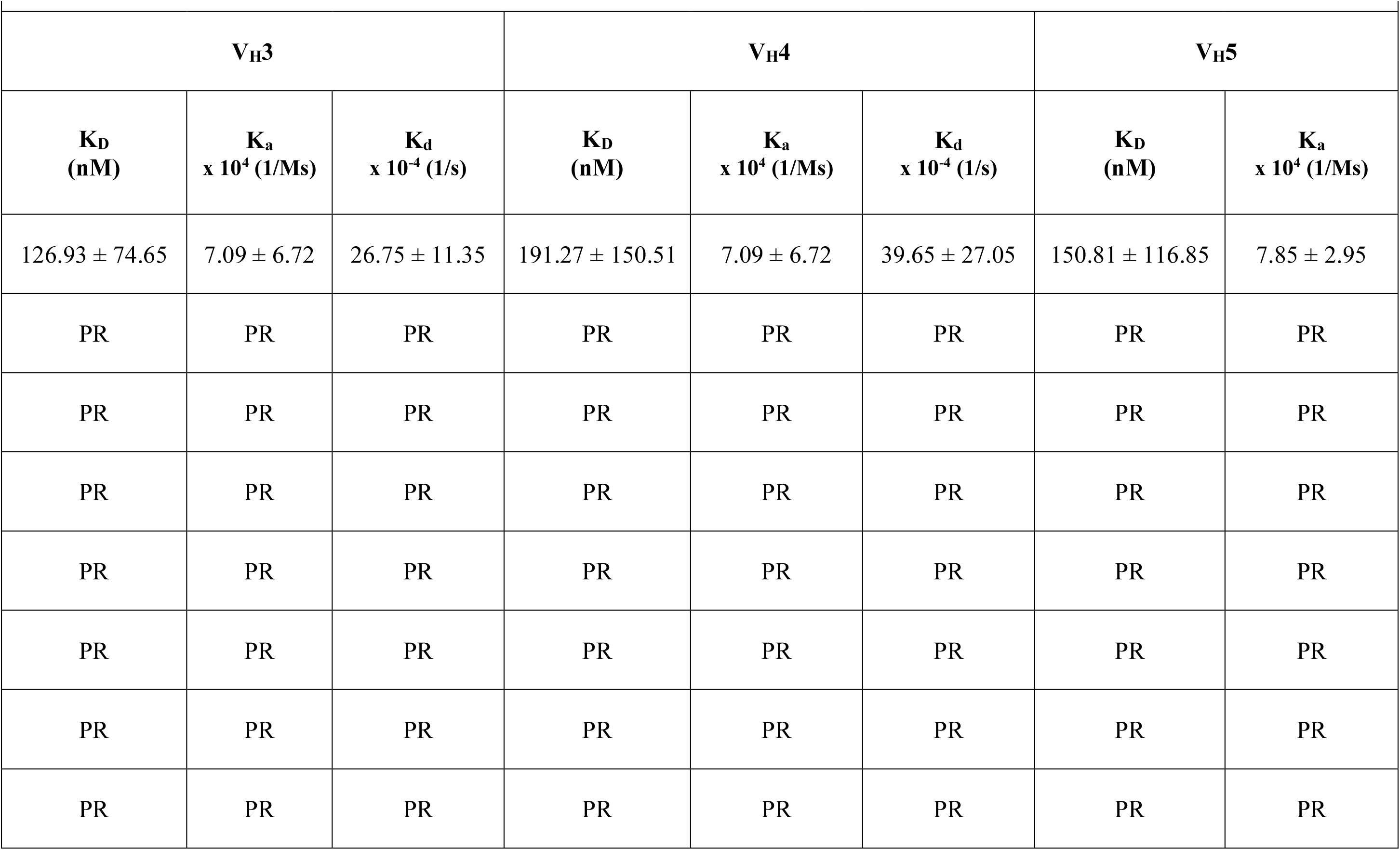
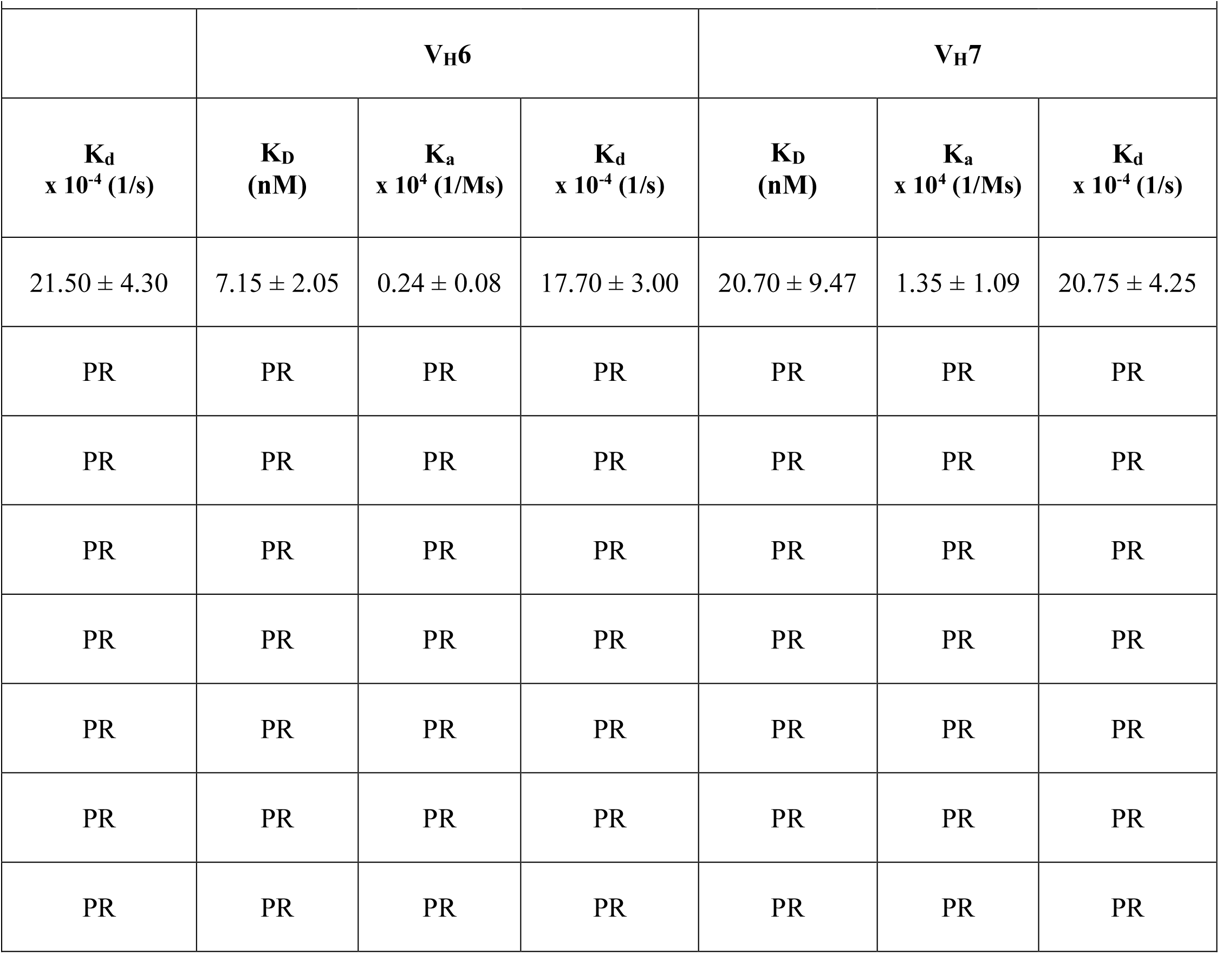
BLI measurement of Pertuzumab IgG_4_ V_H_1 to V_H_7 variants against immobilized FcγRs (FcγRIa, FcγRIIa_H167_, FcγRIIa_R167_, FcγRIIb/c, FcγRIIIa_F176_, FcγRIIIa_V176_, FcγRIIIb_NA1_ and FcγRIIIb_NA2_). K_D_ (nM), K_a_ (x 10^4^ (1/Ms)) and K_d_ (x 10^-4^ (1/s)) are all presented. PR indicates a poor response (RU below 0.3). All experiments were performed in triplicates with the standard errors shown.

## RESULTS

### IgG_2_ V_H_1-7 – FcγR Binding

The data for IgG_2_ binding are shown in Table 1. The binding between IgG_2_ and FcγRIa was measurable only for V_H_1,3 and 5 families. Each of the three V_H_ family representatives showed similar K_a_ (4.06 to 8.92 × 10^4^ (1/Ms)), K_d_ (5.49 to 15.57 × 10^−4^ (1/s)) and K_D_ (40.70 to 64.00 nM) values. The binding of V_H_2, 4, 6 and 7 IgG_2_ gave a poor response (below RU 0.3).

IgG_2_ was found to have the highest binding affinity for FcγRIIa_H167_, with all V_H_ family representative variants having measurable interactions. Their K_a_ values are very similar (4.86 to 14.02 × 10^4^ (1/Ms)), with V_H_4 being an outlier with a K_a_ of 36.05 × 10^4^ (1/Ms)). Their K_d_ values can be divided into two groups: those with higher dissociation rates, such as for V_H_1, 2, 3 and 7 (4.86 to 9.76 × 10^−4^ (1/s)) and those with weaker dissociation rates, as seen for V_H_4, 5 and 6 (14.87 to 21.14 × 10^−4^ (1/s)). The K_D_ values was also between the range of 9.58 to 33.30 nM). IgG_2_ interaction with FcγRIIa_R167_ was highly dependent on the V_H_ representative family, with V_H_1, 2, 3 and 7 showing poor binding, whereas V_H_4, 5 and 6 showed similar K_a_ (between 1.33 and 1.83 × 10^4^ (1/Ms)), K_d_ (between 5.94 and 7.71 × 10^−4^ (1/s)) and K_D_ (between 18.00 and 26.05 nM).

Among the receptors, FcγRIIb/c had the weakest K_D_ interaction with IgG_2,_ with V_H_2 and V_H_7 showing poor responses. The K_a_ (between 5.17 and 6.09 × 10^4^ (1/Ms)) of V_H_3,5 and 6 IgG_2_ was similar. V_H_1 and V_H_3 showed weaker association rates (17.56 and 30.75 × 10^4^ (1/Ms), respectively). The K_d_ (between 3.97 and 7.62 × 10^−4^ (1/s) for all binding IgG_2_s) had the V_H_4 representative variant showing a noticeably weaker dissociation (93.37 × 10^−4^ (1/s)) than the other family representatives. The K_D_ values of the IgG2-FcγRIIb/c were weak between 47.75 and 355.00 nM.

IgG_2_ interactions with the FcγRIII variants (FcγRIIIa_F176, V176_, FcγRIIIb_NA1_ and _NA2_ in Table 1) all gave poor responses regardless of the V_H_ family. Consistent with our previous work on IgG_1_, our V_H_2 FWR grafted IgG_2_ generally gave poor responses. Notably, V_H_7 IgG_2_ had better (smaller) K_D_ values, and V_H_1, 3, 4, 5 and 6 family variants showed similar binding affinity when compared to one another for binding FcγRs in Table 1.

### IgG_3_ V_H_1-7 – FcγR Binding

Table 2 shows IgG_3_ interacted strongly with FcγRIa regardless of the V_H_ family used, except for the V_H_3 representative variant, which resulted in a poor response. Except for VH1, with a weaker association rate of 20.76 × 10^4^ (1/Ms)), the K_a_ was very similar (2.46 and 4.19 × 10^4^ (1/Ms)). The K_d_ was closer between V_H_2, 4, 5, 6 and 7 (between 0.64 and 2.01 × 10^−4^ (1/s)), with V_H_1 showing a weaker dissociation rate of 5.53 × 10^−4^ (1/s). The K_D_ for all variants, except for VH3 having poor responses, were between 0.63 and 2.90 nM.

For FcγRIIa_H167_, the K_a_ of the VH IgG_3_ variants were between 7.46 to 13.21 × 10^4^ (1/Ms)). The K_d_ of these variants were within a similar range of strength (between 3.95 and 7.11 × 10^−4^ (1/s)), with VH5 being an outlier at 27.85 × 10^−4^ (1/s). Regarding FcγRIIa_H167_, the IgG_3_ K_D_ was also identified to be very similar (between 14.97 and 23.56 nM), with VH3 having a much stronger K_D_ (5.75 nM) and V_H_4 displaying a poor binding response.

Regarding FcγRIIa_R167_, all the IgG_3_s did not yield a measurable response.

All V_H_ chains IgG_3_s were able to bind FcγRIIb/c. The K_a_ of the variants to the receptor was found to be very similar, being between 1.50 and 2.67 × 10^4^ (1/Ms), with V_H_4 displaying a slightly weaker association rate of 7.29 × 10^4^ (1/Ms). Their K_d_ values of between 6.28 and 11.01 × 10^−4^ (1/s were all very similar, as were the K_D_ values (13.60 to 16.40 nM), with V_H_5 showing slightly weaker binding affinity with a K_D_ of 24.80 nM.

Only IgG_3_ V_H_1 showed measurable binding to FcγRIIIa_F176_. With a K_a_ of 16.03 × 10^4^ (1/Ms), K_d_ of 328.50 × 10^−4^ (1/s) and K_D_ of 92.90 nM, the rest of the variants showed poor response.

All IgG_3_s could bind FcγRIIIa_V176_, with the K_a_ values being similar in a narrow range between 3.48 and 4.94 × 10^4^ (1/Ms), with VH3 displaying a stronger association rate of 1.35 × 10^4^ (1/Ms). The K_d_ values were also very similar and in a narrow range between 8.12 and 9.89 × 10^−4^ (1/s)) except for the V_H_3 IgG_3,_ which had a significantly weaker dissociation rate of 177.77 × 10^−4^ (1/s). The K_D_ values were also similar for V_H_1, 2, 5, 6 and 7, between 13.99 and 28.60 nM) with V_H_3 and V_H_4 IgG_3_s having 181.18 and 88.67 nM,.

IgG_3_ V_H_1, 2, 3, 4 and 5 bound to FcγRIIIb_NA1_, with the exception of V_H_6 and 7 showing poor responses. The K_a_ of V_H_1, 2, 3, 4 and 5 were in a narrow range between 0.22 and 0.71 × 10^4^ (1/Ms), as were the K_d_ values (between 121.50 and 188.67 × 10^−4^ (1/s)). The K_D_ values were thus expectedly similar (between 17.29 and 52.10 nM).

For FcγRIIIb_NA2,_ only IgG_3_ V_H_3 showed any measurable binding with the K_a_ of 1.51 × 10^4^ (1/Ms)), the K_d_ at 28.05 × 10^−4^ (1/s), and the K_D_ at 14.58 nM. The rest of the IgG_3_s did not show measurable responses.

### IgG_4_ V_H_1-7 – FcγR Binding

In measuring our IgG_4_ panel, only FcγRIa showed any readings using BLI (Table 3). All other FcγRs (FcγRIIa_H167_, FcγRIIa_R167_, FcγRIIb/c, FcγRIIa_F176_, FcγRIIIb_NA1_ and FcγRIIIb_NA2_) showed a poor response.

With all the IgG_4_ V_H_ variants binding to FcγRIa, The Ka values were between d 7.85 × 10^4^ (1/Ms). The K_d_ were also in the narrow range between 17.25 and 39.65 × 10^−4^ (1/s). The K_D_ values could be divided into three distinct groups, strong binders: V_H_1 and V_H_6 (3.62 and 7.15 nM, respectively), intermediate binders: V_H_2 and V_H_7 (75.72 and 20.70 nM, respectively) and weak binders: V_H_3, 4 and 5 (126.93, 191.27 and 150.81 nM respectively).

### BLI measurements of immobilized bacterial B-cell superantigens Protein G and Protein L to Pertuzumab V_H_1-V_H_7 IgG_2, 3_ and _4_ variants

The binding affinity of V_H_1-V_H_7 IgG_2, 3_ and _4_ Pertuzumab variants to Protein L (PpL) and Protein G (SpG) were similar and within the narrow ranges of 0.29 nM to 6.17 nM and 0.80 nM to 4.19 nM, respectively (Table 4).

**Table 4:**
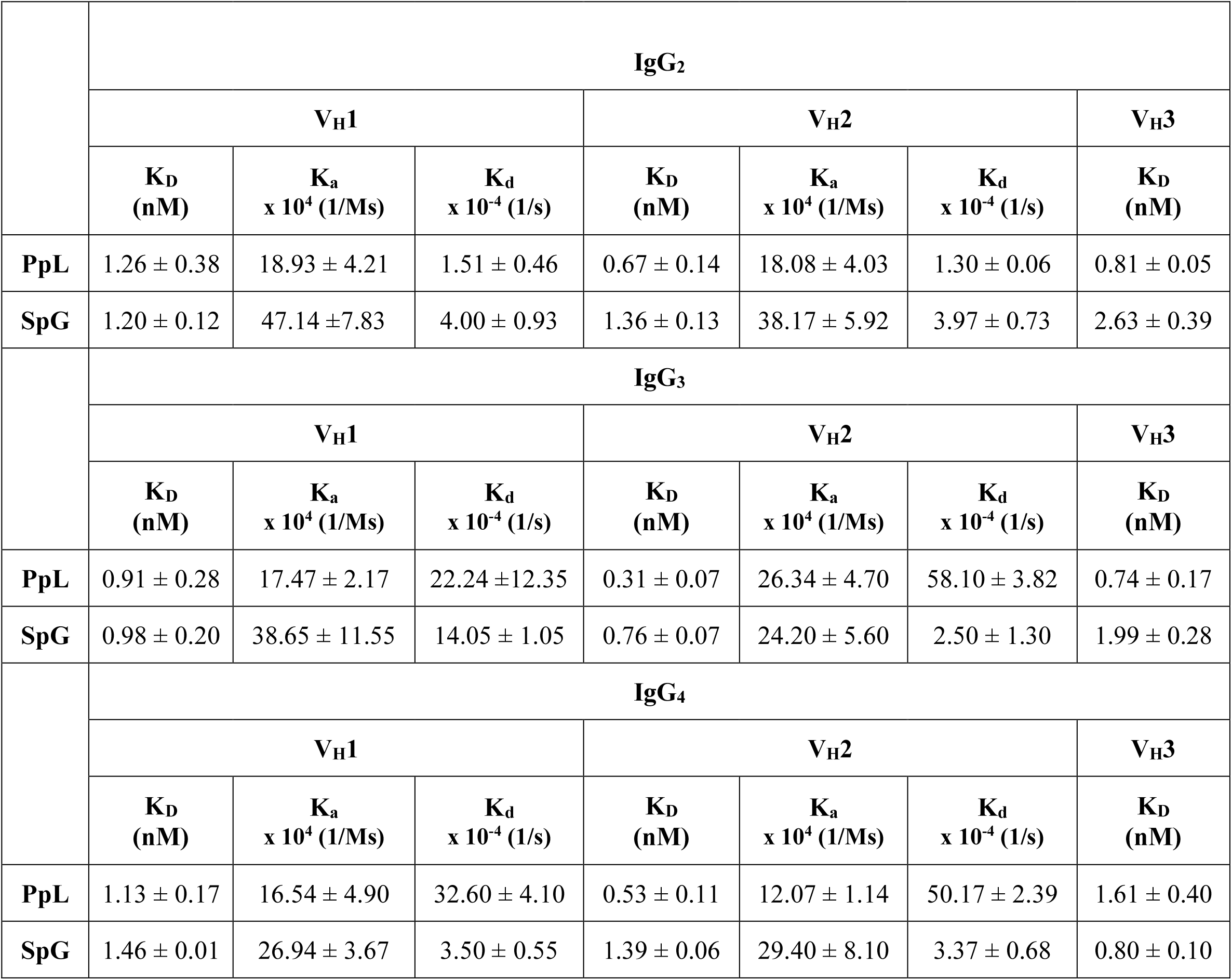
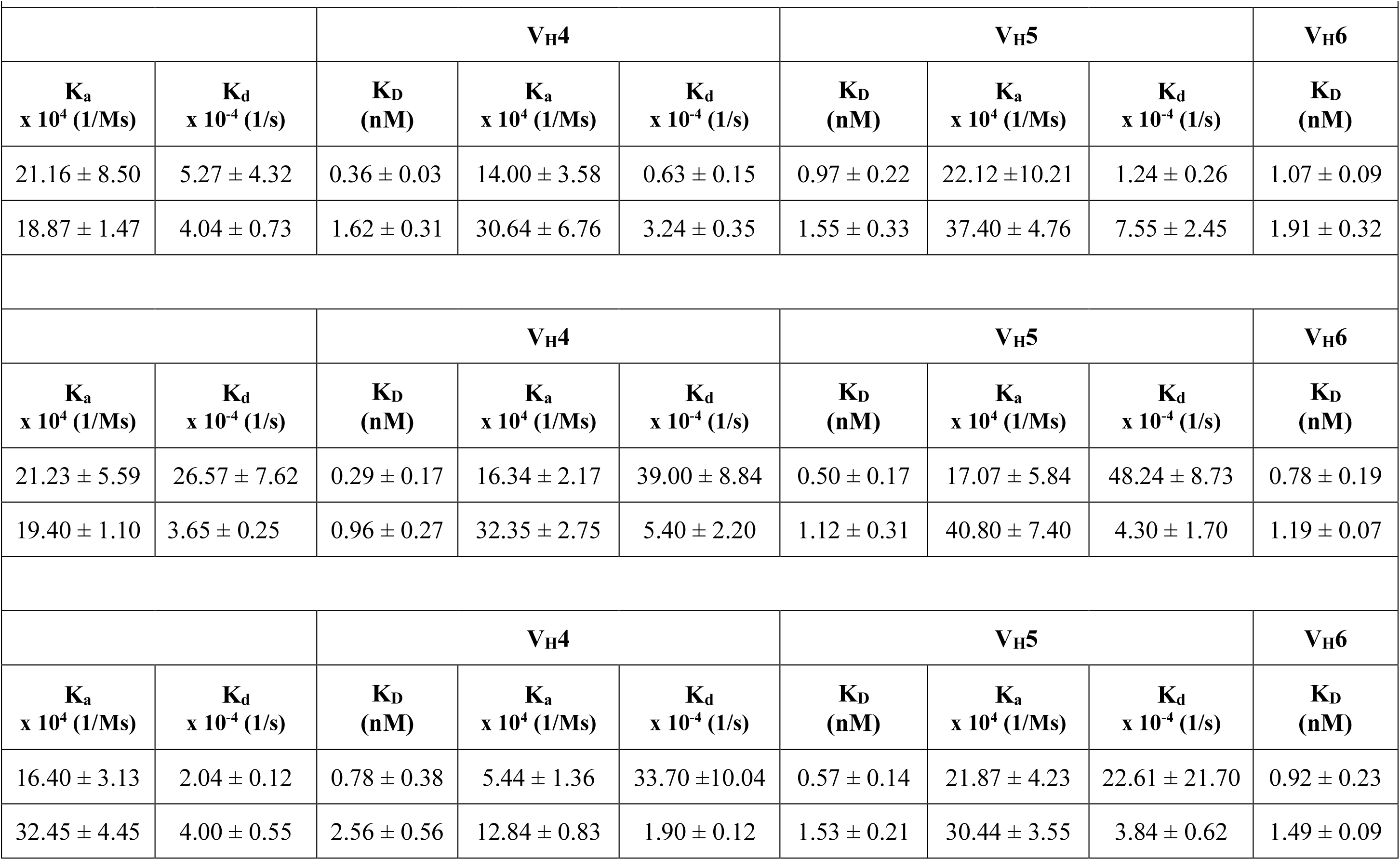
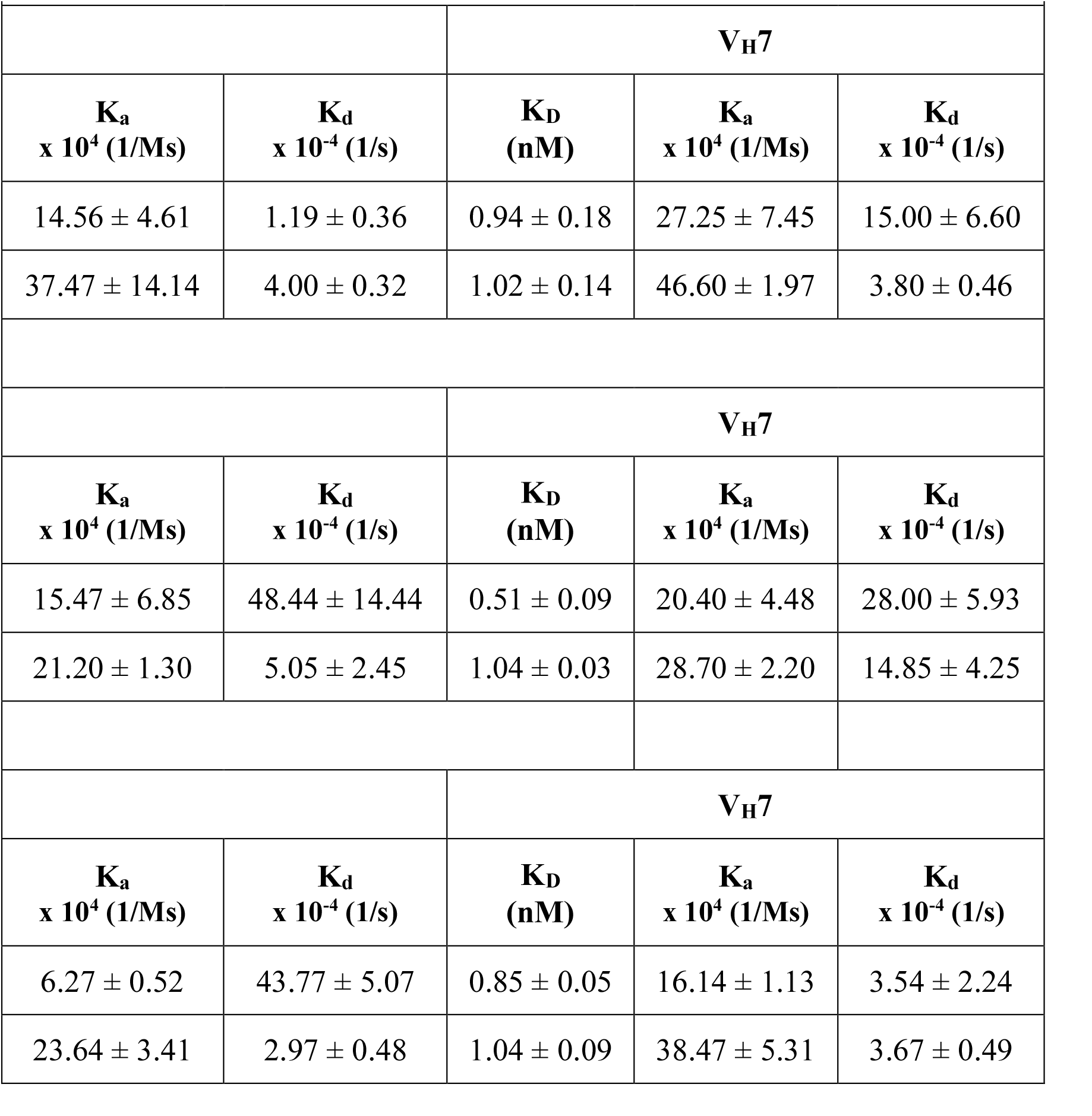
BLI measurement of Pertuzumab IgG_2, 3 and 4_ V_H_1 to V_H_7 variants against immobilized Protein L (PpL) and Protein G (SpG). K_D_ (nM), K_a_ (x 10^4^ (1/Ms)) and K_d_ (×10^-4^ (1/s)) are all presented. PR indicates a poor response (RU below 0.3). All experiments were performed in triplicates with the standard errors shown.

### Protein L – IgG_2,3_ and _4_ Binding

All IgG_2_,_3_ and _4_ V_H_ variants with the same Vκ1 showed measurable PpL binding to the IgG_2_ variants (Table 4).

For IgG_2_, the K_a_ values were within a narrow range between 14.00 and 27.25 × 10^4^ (1/Ms). The K_d_ values were more diverse between 0.63 and 15.00 × 10^−4^ (1/s). Despite the significant variation in the K_d_, the K_D_ values were within a narrow range between 0.36 and 1.26 nM).

For IgG_3_ binding to PpL (Table 4), the K_a_ values fall within a similarly narrow range of between 15.47 and 26.34 × 10^4^ (1/Ms), as were the K_d_ values being between 22.24 and 58.10 × 10^−4^ (1/s), to result in an expected narrow range for the K_D_ values between 0.29 and 0.91 nM.

The binding of the IgG_4_ V_H_ variants to PpL (Table 4) showed the K_a_ values were between 5.44 and 21.87 × 10^4^ (1/Ms), with the K_d_ values between 2.04 and 50.17 × 10^−4^ (1/s) to result in K_D_ values between 0.53 and 1.61 nM.

Collectively, PpL bound the IgG_2_,_3_ and _4_ V_H_ variants within a relatively narrow range between 0.29 to 1.61 nM.

### Protein G – IgG_2,3_ and _4_ Binding

Shown in Table 4, all IgG_2_,_3_ and _4_ V_H_ variants bound Protein G (SpG). IgG_2 –_ SpG showed a K_a_ range of 18.87 to 47.14 × 10^4^ (1/Ms) and a narrow K_d_ range of 3.24 and 7.55 × 10^−4^ (1/s) to have the calculated narrow K_D_ range between 1.02 and 2.63 nM.

The IgG_3_ – SpG panel showed a narrow K_a_ range of 19.40 and 47.14 × 10^4^ (1/Ms) and a K_d_ range between 2.50 and 14.85 × 10^−4^ (1/s) to yield a K_D_ range between 1.02 and 2.63 nM.

The IgG_4_ – SpG panel showed a K_a_ range of 12.84 and 38.47 × 10^4^ (1/Ms), a narrow K_d_ range between 1.90 and 3.84 × 10^−4^ (1/s) to yield K_D_ values 0.80 and 2.56 nM.

Collectively, SpG bound the IgG_2_,_3_ and _4_ V_H_ variants within a relatively narrow range between 0.80 to 2.63 nM.

## DISCUSSION

Combining our previously V_H_1-7 grafted Trastuzumab IgGs [23] with our Pertuzumab IgG_2_,_3_ and _4_ [3], we generated V_H_1-7 variants of IgG_2_,_3_ and _4_s. This panel completes and further expands on our previous work on IgG_1_ regarding FcR binding [23] and other antibody isotypes of IgA_1_, IgA_2_, IgE and IgM [5, 7, 23] with their respective FcRs to form a more comprehensive investigation. Through the created representative V_H_ family 1-7 variants of IgG_2_,_3_ and _4_s with the Pertuzumab CDRs and Vκ1 light chain, this work experimentally measured using BLI, the IgGs interaction with FcγRIa, FcγRIIa_H167,_ FcγRIIa_R167_, FcγRIIb/c, FcγRIIIa_F176_, FcγRIIIa_V176_, FcγRIIIb_NA1_ and FcγRIIIb_NA2_.

The BLI results with the FcγRs showed that the V_H_ family FWRs significantly influenced receptor engagement. While IgG_2_ was previously reported to be unlike other IgG isotypes to not bind to FcγRIa [1] due to the loss of the L235 [28], our IgG_2_s, when grafted with V_H_1, 3, and V_H_5 could. IgG_4_, on the other hand, while able to bind FcγRIa, did so at a lower affinity than IgG_3_ [15]. Our IgG_4_ variants agree with previous literature [15] that it binds poorly to all other FcγRs.

FcγRIIa is divided into two variants: FcγRIIa_H167_ (low-responder) and FcγRIIa_R167_ (high-responder). FcγRIIa_H167_ has more efficient binding to all IgG isotypes than FcγRIIa_R167_, with the binding strength reported as IgG_3_ > IgG_1_ > IgG_4_ = IgG_2_ in previous work [24]. On the contrary, this investigation found IgG_2_ to have a higher affinity for FcγRIIa_H167_ than FcγRIIa_R167_. IgG_2_ V_H_1, 2, 3 and 7 displayed poor responses to FcγRIIa_R167_ with binding only present for V_H_4, 5 and 6. In our panel, IgG_3_ showed a similar binding affinity towards FcγRIIa_H167_ as IgG_2_, with V_H_4 resulting in a poor response. Our IgG_3_ variants also demonstrated poor binding to FcγRIIa_R167,_ and our IgG_4_ variants had no measurable interactions with both FcγRIIa_H167 and_ FcγRIIa_R167_.

Binding to FcγRIIb/c was generally poor for our tested subtypes compared to previous findings reporting binding affinities of strongest to weakest being IgG_3_ = IgG_1_ = IgG_4_ > IgG_2_ [15]. Our IgG_3_ variants had the highest affinity for FcγRIIb/c with no significant differences between the V_H_ variants. In relative consistency with the previous finding of IgG_2_ being the weakest binder to FcγRIIb/c, our IgG_2_ and IgG_4_ variants generally had poor responses with FcγRIIb/c.

FcγRIIIa is also divided into two variants: FcγRIIIa_F176_ and FcγRIIIa_V176_. Both FcγRIIIa variants have a high affinity for each IgG isotype (IgG_3_ > IgG_1_ > IgG_4_ > IgG_2_), as previously reported [1]. Our IgG_2_ generally showed a poor response to both FcγRIIIa_F176_ and FcγRIIIa_V17,6,_ whereas apart from our IgG_3_ VH3 with lower binding, the rest of the IgG_3_ variants showed strong binding to FcγRIIIa_V176_. IgG_3_ binding to FcγRIIIa_F176_ shows poor binding except for V_H_1, which displayed weak binding. IgG_4_ had a poor binding response to FcγRIIIa_F176_ and FcγRIIIa_V176_.

The final FcγR measured is tested as FcγRIIIb_NA1_ and FcγRIIIb_NA2_. Both FcγRIIIb variants are known to bind IgG_3_ but not to IgG_2_ or IgG_4_ [15]. This investigation reflects this for IgG_2_ and IgG_4,_ as both showed no measurable responses regardless of the V_H_ sequence present. IgG_3_ was measured to bind all FcγRIIIb_NA1_ except for V_H_6 and V_H_7 resulting in poor binding. IgG_3_ could not bind FcγRIIIb_NA2_, although V_H_3 grafted IgG_3_ showed strong binding affinity. This thus suggests that previously reported interactions between IgG_3_ and FcγRIIIb_NA_ did not apply to all IgG3s.

B-cell superantigens bind antibodies outside the CDRs to be non-specific binders [29]. PpL and SpG are commonly studied B-cell superantigens with PpL canonically binding to the Vκ1, 3 and 4 light chains [30, 31] with exceptions while SpG canonically binds to the C_H_2 – C_H_3 cleavage site on the IgG Fc [32]. By grafting different V_H_ regions, we showed that PpL continued to bind the κ light chain with minimal variation. SpG also had a narrow fluctuation in binding to the Fc region of IgG_2, 3_ and _4_. Such minor variations for PpL and SpG show the robustness of these microbial B-cell superantigens that were more independent of V_H_ influences than the FcγRs. These findings agree with and supplement previous research on V_H_ and κ chain variation on IgG_1_ -Protein L / Protein A binding [20].

The findings here with the PpL and SpG demonstrate the robust functionality of these microbial superantigens in immune evasions, even beyond the distal allosteric effects from the V-regions. As such, it supports that the use of these superantigens in antibody purification remains reliable, as are biophysical measurement methods that capture antibodies using these superantigens. While the effect on antigen and FcR binding by the prior binding of these superantigens on the antibodies remains to be fully elucidated mechanistically, it would undoubtedly be required to characterise future biologics given their potential presence among normal flora.

While binding affinity is not directly correlated to the efficiency of effector function activation [33], we noted extreme non-canonical interactions of the IgGs with their receptors, where previously reported non-binders could, and reported binders, could not. Such extreme non-canonical bindings would certainly drastically affect the function of such IgG therapeutics that may explain the often unreported loss or gain of effects during humanization. With ethnic polymorphisms present in the FcγRs[34], it is essential to understand how changing the V_H_ sequence can elicit these mutations to influence Fc – receptor binding and its downstream effects on FcγR signalling. IgG_1_ and IgG_3_ are the most potent antibodies [1] in therapeutics. This is supported by our findings here regarding IgG_3_ further showed that it can bind the tested FcγRs at a higher affinity than the subtypes, with a less pronounced effect from the distal efforts of V_H_-FWRs, with particular attention to the VH_3_-FWR.

In conclusion, our findings show the significant influence from the V_H_-FWRs on IgG_2,3_ and _4_ engagements of FcγRs, at times to an extreme manner. Such interactions are likely to cause dramatic functional loss or gain of antibody therapeutics that need to be considered during humanization and characterization. While the lack of drastic effects in PpL and SpG could indicate their reliability in the use for antibody purification and capture in certain assays, care should also be noted in their possible interaction during clinical use if such antibody superantigens would be present in the normal flora. Together, this report challenged previous canonical reported interactions that can serve as a reminder on the need for detailed investigation on any IgG in development with regards to the assumed expected functions therapeutically and other applications.

## AUTHOR CONTRIBUTIONS

AD performed the BLI experiments with the receptors and the superantigens, as well as drafted the manuscript. SKEG conceived the project, provided the resources for the experiments, supervised the entire project and also revised and edited the manuscript.

## FUNDING

AD is a PhD student funded by The University of Manchester, UK and the Agency for Science, Technology and Research (A*STAR), Singapore. This work was partially supported by the Wenzhou Science and Technology Bureau, Key Lab Program, Wenzhou Municipal Key Laboratory for Applied Biomedical and Biopharmaceutical Informatics, Wenke Jiji (2021) No. 4, to Wenzhou-Kean University.

## CONFLICT-OF-INTEREST

All authors declared no conflict of interest.

## ETHICS AND CONSENT STATEMENT

No animals were used in this work.

## ACKNOWLEDGEMENTS

We would like to to thank JYY for helping to express and purify the IgGs.

## DATA AVAILABILITY

Data is available in the Supplementary Material

## Notes

### Competing Interest Statement

The authors have declared no competing interest.

